# HOIL1 regulates group 2 innate lymphoid cell numbers and type 2 inflammation in the small intestine

**DOI:** 10.1101/2022.03.09.483650

**Authors:** Matthew J. Wood, Jeffrey N. Marshall, Victoria L. Hartley, Ta-Chiang Liu, Kazuhiro Iwai, Thaddeus S. Stappenbeck, Donna A. MacDuff

## Abstract

Patients with mutations in *HOIL1* experience a complex immune disorder including intestinal inflammation. To investigate the role of *HOIL1* in regulating intestinal inflammation, we employed a mouse model of HOIL1 deficiency. The ileum of *Hoil1*^−/−^ mice displayed features of type 2 inflammation including tuft cell and goblet cell hyperplasia, and elevated expression of *Il13*, *Il5* and *Il25* mRNA. Inflammation persisted in the absence of T and B cells, and bone marrow chimeric mice revealed a requirement for HOIL1 expression in radiation-resistant cells to regulate inflammation. Although disruption of IL-4 receptor alpha (IL4Rα) signaling on intestinal epithelial cells ameliorated tuft and goblet cell hyperplasia, expression of *Il5* and *Il13* mRNA remained elevated. KLRG1^hi^ CD90^lo^ group 2 innate lymphoid cell (ILC2) were increased independent of IL4Rα signaling, tuft cell hyperplasia and IL-25 induction. Antibiotic treatment dampened intestinal inflammation indicating commensal microbes as a contributing factor. We have identified a key role for HOIL1, a component of the Linear Ubiquitin Chain Assembly Complex, in regulating type 2 inflammation in the small intestine. Understanding the mechanism by which HOIL1 regulates type 2 inflammation will advance our understanding of intestinal homeostasis and inflammatory disorders and may lead to the identification of new targets for treatment.

## Introduction

Inflammatory bowel diseases (IBD) affect around 1% of the US population and prevalence continues to increase in developed countries^1^. IBD is a complex disease influenced by both genetic and environmental factors, and specific treatments are therefore only effective for a subset of patients. Patients with mutations in heme-oxidized IRP2 ubiquitin ligase 1 (*HOIL1*; official gene name *RBCK1*), experience a complex immune disorder involving autoinflammation and inflammatory bowel disease-like symptoms, increased susceptibility to bacterial infections, progressive muscular amylopectinosis and myopathy^2^. Gastrointestinal symptoms in HOIL1-deficient patients include abdominal pain, bloody and mucous stools, colonic lesions and eosinophilic infiltration in the gut epithelium. HOIL1, HOIL1-interacting protein (HOIP; official name RNF31) and SHANK-associated RH domain-interacting protein (SHARPIN) form an E3 ubiquitin ligase complex called the linear ubiquitin chain assembly complex (LUBAC). Patients with mutations in *HOIP* display similar clinical and cellular phenotypes to HOIL1 deficient patients^3^.

LUBAC is the only enzyme known to generate linear (methionine-1 linked) polyubiquitin chains due to the unique E3 ubiquitin ligase activity of HOIP, and has been shown to regulate NFκB activation and programmed cell death downstream of many innate immune receptors, including TNFR1, IL1R1, IL-17R and toll-like receptors (TLRs)^4–6^. LUBAC also regulates CD40, B and T cell receptor, inflammasome and RIG-I-like receptor signaling pathways. Accordingly, HOIL1 and LUBAC are important for the efficient induction of type 1 inflammatory cytokines and interferons, and to control bacterial and viral infections^7–9^.

In mice, complete loss of HOIP or HOIL1 expression results in embryonic lethality due to essential roles in hematopoiesis and in limiting TNFα-induced cell death^10,11^. SHARPIN-deficient mice are viable, but exhibit defects in immune development as well as severe systemic inflammation within the first two months of life^12^. To study the physiological consequences of OIL1-deficiency, we have employed a HOIL1-mutant mouse model (*Hoil1*^−/−^ herein) that expresses the N-terminal domain of HOIL1 at approximately ten percent of wild-type levels, enabling partial stabilization of LUBAC and viability of homozygous mice^8,13,14^. Expression of both HOIL1 and HOIP is reduced, and LUBAC function is impaired in mouse embryonic fibroblasts and bone marrow-derived macrophages from these mice^8,13,14^. We previously demonstrated that these mice are a relevant model of human HOIL1-deficiency, since they exhibit immunodeficiency or hyperinflammatory responses, depending on the pathogenic challenge^8^. Macroscopically, naïve *Hoil1*^−/−^ mice are largely indistinguishable from their wild-type (*Hoil1*^+/+^) littermates, but glycogen-like deposits are observed in the cardiac tissue of mice by 18 months of age, similar to those observed in humans with mutations in HOIL1^8^.

Here, we show that expression of HOIL1 in a radiation-resistant cell type is required to limit type 2 inflammation in the small intestine. Excessive expression of type 2 inflammatory cytokines in HOIL1-deficient tissue did not require T cells or B cells, tuft cell hyperplasia or IL4Rα-dependent induction of IL-25. Global gene expression and flow cytometric analyses indicated that group 2 innate lymphoid cell (ILC2) numbers were increased in the absence of HOIL1 and independent of IL-25 induction. Antibiotics treatment alleviated the inflammation, indicating a role for microbial sensing. Our data reveal a novel role for HOIL1 in regulating type 2 inflammation in the intestine, contributing to a broader understanding of the mechanisms of intestinal homeostasis and disease.

## Results

### HOIL1-deficient mice exhibit type 2 intestinal inflammation in the distal ileum

To investigate whether HOIL1 deficiency causes intestinal inflammation in mice, we examined the distal ileum from specific pathogen-free (SPF) *Hoil1^+/+^* and *Hoil1*^−/−^ co-housed littermates. Histological analysis revealed goblet cell hyperplasia in the distal ileum of *Hoil1*^−/−^ mice (Fig. 1A-C), which was absent in tissues from *Hoil1^+/+^* littermates. This histological change is characteristic of type 2 inflammation observed after intestinal helminth infection^15,16^. Consistently, mRNAs for type 2 inflammatory cytokines *Il4*, *Il5* and *Il13* were elevated in *Hoil1*^−/−^ compared to *Hoil1^+/+^* ileum (Fig. 1D). An increase in IL-4 and IL-5 protein was also detected in the homogenized tissue (Fig. 1E). However, we did not detect changes in type 1 inflammatory cytokine mRNAs, *Ifng* and *Tnf* (Fig. 1F), indicating that *Hoil1*^−/−^ mice do not experience a generalized, non-specific inflammation, or a shift from type 1 to type 2 cytokine production. Type 2 inflammation can be caused by infection with a common intestinal protozoan, *Tritrichomonas muris*, present in some SPF mouse colonies^17^. However, *Tritrichomonas muris* was not detected in fecal samples from our mice (not shown).

**Figure 1:**
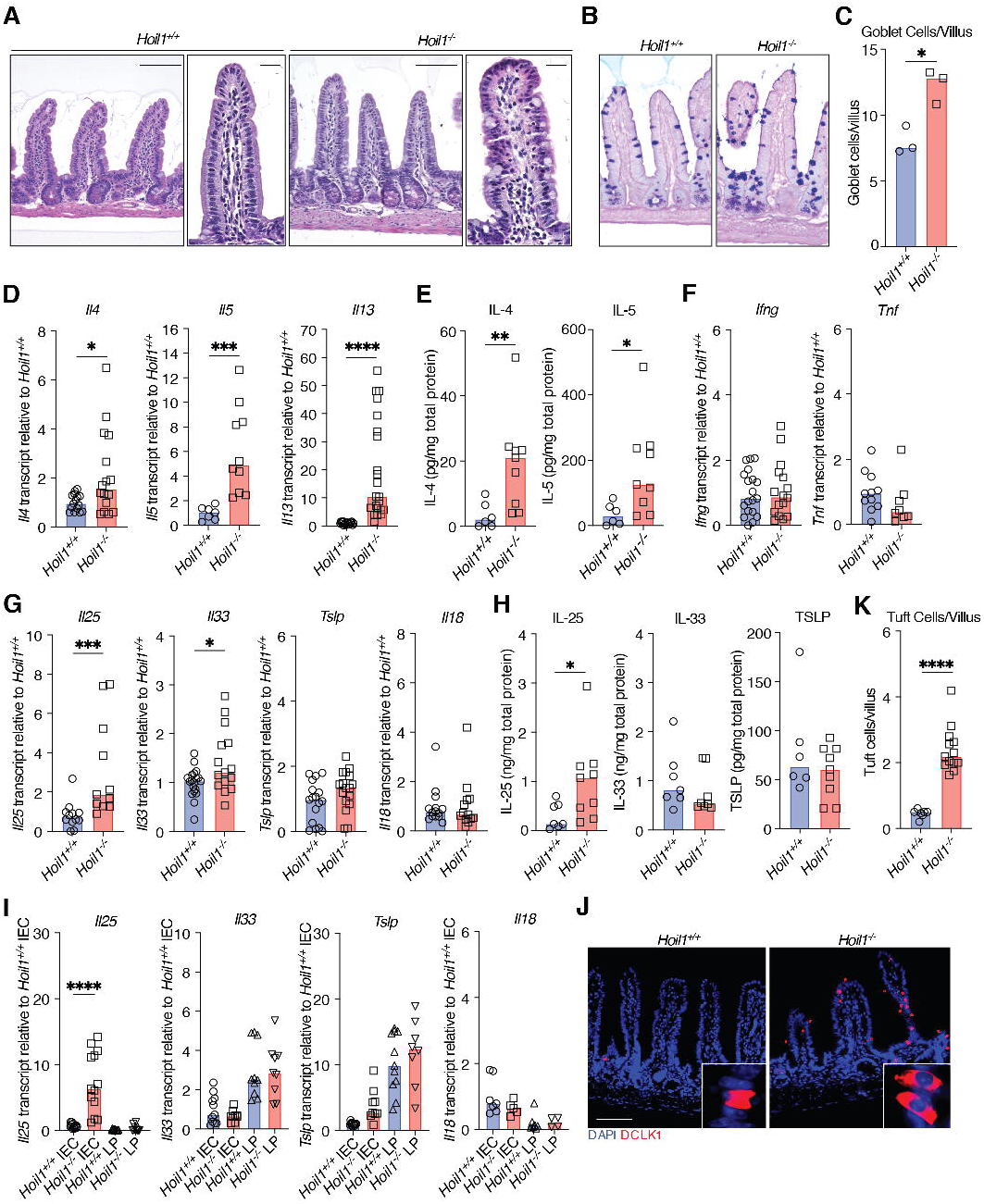
*Hoil1*^−/−^ mice exhibit type 2 inflammation in the distal ileum. (**A**) Representative images of H&E stained sections of ileum from *Hoil1*^+/+^ and *Hoil1*^−/−^ mice. Scale bars represent 50 μm (left panel) and 20 μm (right panels). (**B**) Representative images of PAS/Alcian Blue stained sections. (**C**) Goblet cells per villus in *Hoil1*^+/+^ and *Hoil1*^−/−^ ileum. (**D-H**) Relative *Il4*, *Il5*, *Il13* (D), *Tnf*, *Ifng* (F), *Il25, Il33*, *Il18* and *Tslp* (G) mRNA levels, and IL-4, IL-5 (E), IL-25, IL-33 and TSLP (H) protein levels in ileum of *Hoil1*^+/+^ and *Hoil1*^−/−^ mice. (**I**) *Il18*, *Il25*, *Il33* and *Tslp* mRNA levels in *Hoil1*^+/+^ and *Hoil1*^−/−^ IEC and LP fractions relative to *Hoil1*^+/+^ IEC median. (**J**) DCLK1 (red) and DAPI (blue) stained sections of ileum from *Hoil1*^+/+^ (left) and *Hoil1*^−/−^ (right) mice (scale = 50 μm). (**K**) Enumeration of tuft cells in villi and crypts of *Hoil1*^+/+^ and *Hoil1*^−/−^ ileum. Each symbol represents a sample from an individual mouse and colored bars represent the median. mRNA levels are expressed as relative to the median level for *Hoil1^+/+^*. Histological enumerations and measurements represent the mean from >10 villi per mouse. All mice were aged between 6-9 weeks. H&E = hematoxylin and eosin; IEC = intestinal epithelial cell fraction; LP = lamina propria fraction. \**p* ≤ 0.05, \*\**p* ≤ 0.01, \*\*\**p* ≤ 0.001, \*\*\*\**p* ≤ 0.001 by Mann-Whitney (C-H, K) or ordinary 2-way ANOVA (I).

Production of IL-4, IL-5 and IL-13 can be induced by IL-25, IL-33, TSLP and IL-18^16,18,19^. *Il25* and *Il33* mRNAs and IL-25 protein were slightly elevated in *Hoil1*^−/−^ ileum relative to *Hoil1*^+/+^ ileum (Fig. 1G, H). However, no differences in *Il18* and *Tslp* mRNA, or in IL-33 and TSLP total protein, were observed. In order to increase the sensitivity of cytokine mRNA detection, we mechanically separated the epithelial cell (IEC) and lamina propria (LP) cell fractions. Expression of *Il33, Tslp* and *Il18* mRNA was similar for *Hoil1^+/+^* and *Hoil1*^−/−^ ileum within each cell fraction (Fig. 1I). However, *Il25* mRNA was significantly higher in the *Hoil1*^−/−^ epithelial cell fraction. Tuft cells are the primary producers of IL-25 in the small intestine, undergo hyperplasia in response to IL-13 during helminth infection, and can be identified by their unique expression of DCLK1^20^. Accordingly, DCLK1^+^ cells with tuft cell morphology were significantly increased in the distal ileum of *Hoil1*^−/−^ mice (Fig. 1J, K). Taken together, these data show that HOIL1 deficiency in mice results in a type 2-like inflammation in the distal ileum associated with excessive expression of *Il4*, *Il5*, *Il13* and *Il25* mRNA and histological changes.

### Symbiotic microbes promote type 2 inflammation in the absence of HOIL1

We next examined the post-natal development of type 2 inflammation in the ileum of *Hoil1*^−/−^ mice. No significant differences in *Il5*, *Il13* or *Ifng* mRNA expression were measured in the intestine from newborn *Hoil1^+/+^* and *Hoil1*^−/−^ mice (Fig. 2A). At 3 weeks of age, both *Il5* and *Il13* mRNAs were slightly, but not significantly, elevated in the ileum of *Hoil1*^−/−^ mice, and at lower levels than observed in 6 to 8 week-old *Hoil1*^−/−^ mice (Fig. 2B, 1D). These data suggest that *Hoil1*^−/−^ mice develop intestinal inflammation with age, possibly due to increasing microbial exposure and diversity. To test whether colonization with intestinal microbes drives this intestinal inflammation in *Hoil1*^−/−^ mice, we treated 6 to 8 week-old mice with a broad-spectrum cocktail of antibiotics for two weeks by daily oral gavage. At 7 and 14 days after starting antibiotics treatment, bacterial 16S DNA levels in stool were below the limit of detection (Fig. 2C). Following 14 days of antibiotic treatment, expression of *Il4*, *Il5* and *Il13* mRNA in distal ileum of *Hoil1*^−/−^ mice was reduced to the level measured in *Hoil1*^+/+^ mice treated with water (Fig. 2D-E). These mRNAs were also reduced in the ileum of *Hoil1^+/+^* mice by antibiotics treatment, but to a lesser extent. No differences in *Il18*, *Tslp*, *Il33, Ifng* or *Tnf* expression were measured between *Hoil1*^+/+^ and *Hoil1*^−/−^ mice, although expression was reduced by antibiotics treatment (Fig. 2E, F). *Il25* remained slightly elevated in tissue from antibiotics-treated *Hoil1*^−/−^ mice despite *Il13* and *Il4* being reduced to water-treated *Hoil1*^+/+^ levels or below. The number of goblet and tuft cells in antibiotics-treated *Hoil1*^−/−^ mice was reduced almost to *Hoil1*^+/+^ frequencies, which may be a direct effect of loss of microbial exposure to the IECs, or an indirect effect via a reduction in IL-13 expression (Fig. 2G-J). These data indicate that microbial exposure contributes to aberrant type 2 inflammation in the absence of HOIL1.

**Figure 2:**
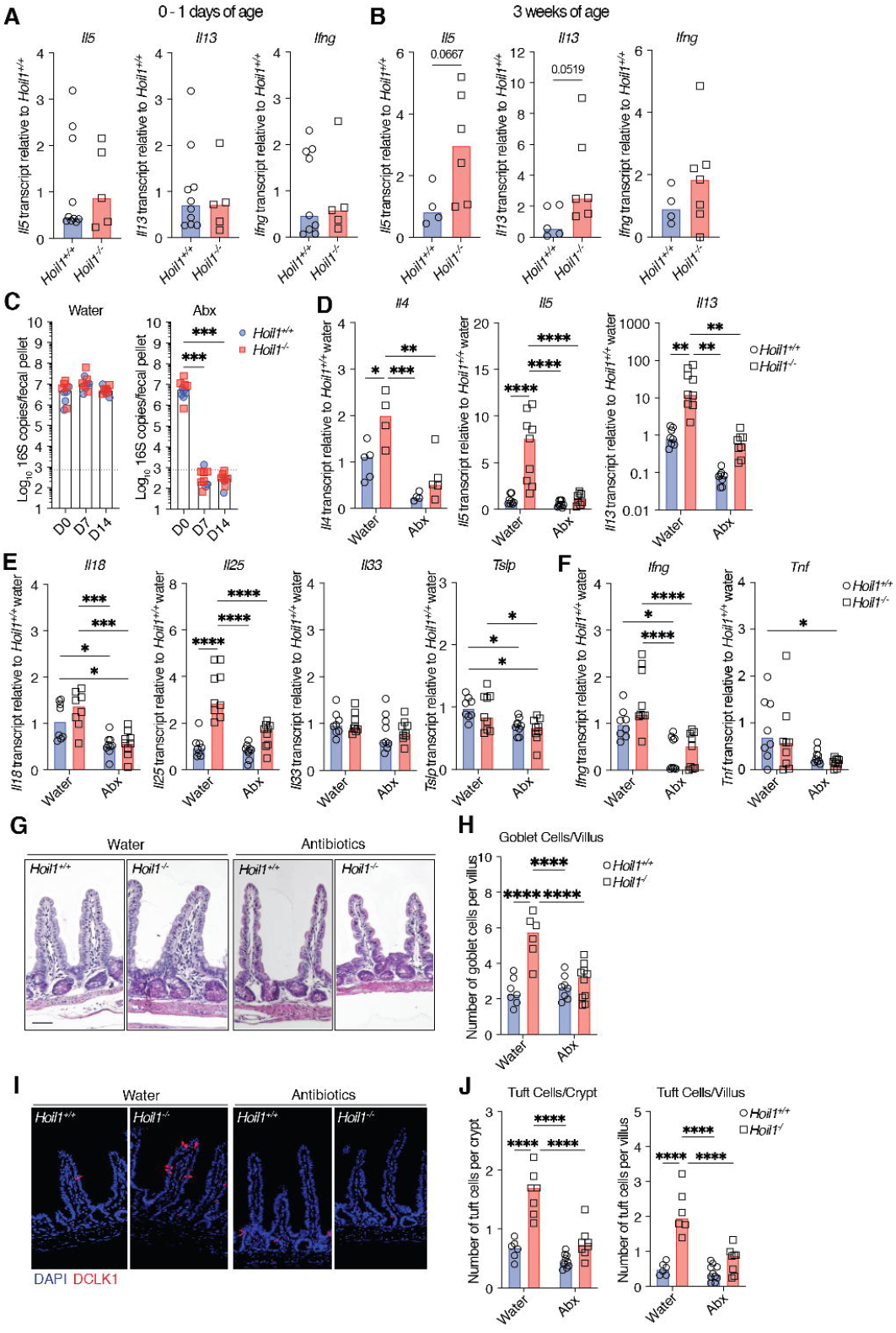
Commensal microbes promote type 2 inflammation in the absence of HOIL1. (**A-B**) *Il5*, *Il13* and *Ifng* mRNA levels in *Hoil1*^+/+^ and *Hoil1*^−/−^ ileum from newborn (**A**) and 3 week-old (**B**) mice. (**C**) 16S rDNA copies per fecal pellet from *Hoil1*^+/+^ or *Hoil1*^−/−^ mice day 0, 7 or 14 after commencement of daily gavage with water (left panel) or antibiotics (vancomycin, neomycin, ampicillin, metronidazole, Abx, right panel). (**D-F**) *Il4, Il5*, *Il13* (**D**), *Il18*, *Il25, Il33*, *Tslp* (**E**), *Ifng* and *Tnf* (**F**) mRNA levels in ileum from *Hoil1*^+/+^ or *Hoil1*^−/−^ mice after 14 days of daily gavage with water or antibiotics (Abx). mRNA levels are relative to *Hoil1*^+/+^. (**G-H**) H&E stained sections (**G**; scale = 50 μm), and enumeration of goblet cells in villi (**H**) in the ileum from *Hoil1*^+/+^ and *Hoil1*^−/−^ mice after 14 days of daily gavage with water or antibiotics. (**I-J**) DCLK1 (red) and DAPI (blue) stained sections (**I**; scale = 50 μm), and enumeration of tuft cells in villi and crypts (**J**) from ileum from *Hoil1*^+/+^ and *Hoil1*^−/−^ mice after 14 days of daily gavage with water or antibiotics. Each symbol represents a sample from an individual mouse and colored bars represent the median. Histological enumerations represent the mean from of >10 villi/crypt per mouse. Mice were aged between 6-9 weeks unless stated otherwise. H&E = Hematoxylin and Eosin. \**p* ≤ 0.05, \*\**p* ≤ 0.01, \*\*\**p* ≤ 0.001, \*\*\*\**p* ≤ 0.001 by Mann-Whitney (A, B), Kruskal-Wallis test with Dunn’s multiple comparisons (C), or ordinary two-way ANOVA with Tukey’s multiple comparisons test (D-F, H, J).

### Excess production of *Il13* and *Il5* occurs independently of goblet and tuft cell hyperplasia and IL-25 induction in *Hoil1*^−/−^ ileum

During helminth infection, IL-13 stimulation of IECs drives epithelial cell changes, including goblet and tuft cell hyperplasia similar to that observed in the *Hoil1*^−/−^ mice. Through a feed-forward mechanism, increased production of IL-25 by tuft cells promotes further production of IL-13, IL-5 and IL-4^16,18,19^. To determine whether the elevated levels of IL-13 were responsible for the epithelial abnormalities observed, we examined the role of IL-13/IL-4 signaling specifically in IECs by crossing *Hoil1*^+/−^ mice to *Il4ra*^flox/flox^ mice and VillinCre (ΔIEC) transgenic mice^21,22^. Histological analysis of the distal ileum from these mice revealed that deletion of IL4Rα on IECs largely rescued the epithelial cell abnormalities in *Hoil1*^−/−^ mice (Fig. 3A-E). Consistently, *Il25* and *Il33* mRNAs and IL-25 protein were reduced in *Hoil1*^−/−^*Il4ra*^*ΔIEC*^ tissue to levels comparable to *Hoil1*^+/+^*IL4ra*^*f/f*^ and *Hoil1*^+/+^*IL4ra*^*ΔIEC*^ tissue (Fig. 3F, G). As before, no differences in *Tslp* mRNA, or IL-33 and TSLP protein levels were detected across the different genotypes. Surprisingly, *Il5* and *Il13* mRNAs remained elevated in *Hoil1*^−/−^*Il4ra*^*ΔIEC*^ tissue, despite *Il25* and *Il33* mRNA being reduced to *Hoil1*^+/+^*IL4ra*^*f/f*^ levels (Fig. 3H). IL-4, IL-5 and IL-13 protein was slightly elevated in *Hoil1*^−/−^*Il4ra*^*f/f*^ tissue, but not in *Hoil1*^−/−^*Il4ra*^*ΔIEC*^ tissue, which could indicate that mRNA and protein levels are uncoupled, or reflect the low sensitivity of these assays for whole tissue homogenates (Fig. 3I). *Il13* and *Il5* mRNAs were also elevated in other regions of the gastrointestinal tract such as the jejunum and, to a lesser extent, the mesenteric lymph nodes (MLN) and colon (Fig. 3J-L). Differences in *Il4* mRNA expression were not detectable in the MLN, suggesting that IL-4 may not be a driving component of this pathway. Together, these data show that increased IL-13/IL-4 signaling in IECs via IL-4Rα triggers goblet and tuft cell hyperplasia and the induction of IL-25 in *Hoil1*-deficient ileum, but that an increase in IL-25, IL-33 or TSLP is not required to drive the excessive *Il13* and *Il5* mRNA expression.

**Figure 3:**
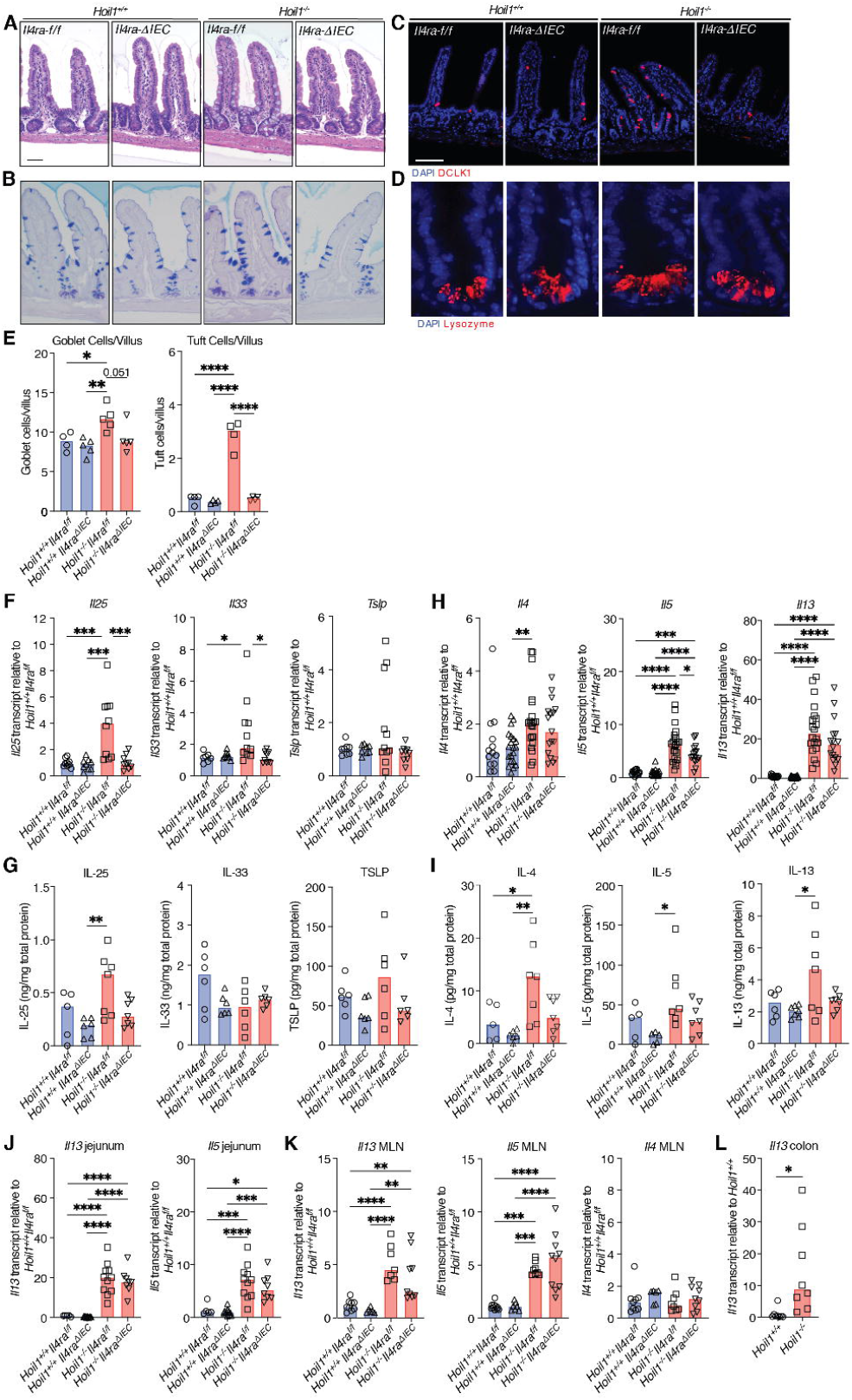
Elevated type 2 inflammatory cytokine expression in *Hoil1*^−/−^ ileum is not dependent on feed-forward signaling in IECs. (**A-D**) H&E (A), PAS/Alcian Blue (B), DCLK1 and DAPI (C), and lysozyme and DAPI (D) stained sections of ileum from *Hoil1*^+/+^*Il4ra*^*f/f*^, *Hoil1*^+/+^*Il4ra*^*ΔIEC*^, *Hoil1*^−/−^*Il4ra*^*f/f*^ and *Hoil1*^−/−^*Il4ra*^*ΔIEC*^ mice (scale = 50 μm). (**E**) Goblet (left panel) and tuft (right panel) cells per villus in *Hoil1*^+/+^*Il4ra*^*f/f*^, *Hoil1*^+/+^*Il4ra*^*ΔIEC*^, *Hoil1*^−/−^*Il4ra*^*f/f*^ and *Hoil1*^−/−^ *Il4ra*^*ΔIEC*^ mice. (**F**) Relative *Il25, Il33* and *Tslp* mRNA levels in ileum from *Hoil1*^+/+^*Il4ra*^*f/f*^, *Hoil1*^+/+^*Il4ra*^*ΔIEC*^, *Hoil1*^−/−^*Il4ra*^*f/f*^ and *Hoil1*^−/−^*Il4ra*^*ΔIEC*^ mice. (**G**) IL-25, IL-33 and TSLP protein levels in distal ileum from *Hoil1*^+/+^*Il4ra*^*f/f*^, *Hoil1*^+/+^*Il4ra*^*ΔIEC*^, *Hoil1*^−/−^*Il4ra*^*f/f*^ and *Hoil1*^−/−^*Il4ra*^*ΔIEC*^ mice. (**H**) Relative *Il4, Il5* and *Il13* mRNA levels in ileum from *Hoil1*^+/+^*Il4ra*^*f/f*^, *Hoil1*^+/+^*Il4ra*^*ΔIEC*^, *Hoil1*^−/−^*Il4ra*^*f/f*^ and *Hoil1*^−/−^*Il4ra*^*ΔIEC*^ mice. (**I**) IL-4, IL-5 and IL-13 protein levels in distal ileum from *Hoil1*^+/+^*Il4ra*^*f/f*^, *Hoil1*^+/+^*Il4ra*^*ΔIEC*^, *Hoil1*^−/−^*Il4ra*^*f/f*^ and *Hoil1*^−/−^*Il4ra*^*ΔIEC*^ mice. (**J-L**) Relative *Il13*, *Il5* and *Il4* mRNA levels in jejunum (**J**), MLN (**K**) and distal colon (**L**) from *Hoil1*^+/+^*Il4ra*^*f/f*^, *Hoil1*^+/+^*Il4ra*^ΔIEC^, *Hoil1*^−/−^ *Il4ra*^*f/f*^ and *Hoil1*^−/−^*Il4ra*^*ΔIEC*^ mice. Each symbol represents a sample from an individual mouse and colored bars represent the median. mRNA levels are expressed as relative to the median for *Hoil1*^+/+^*Il4ra*^*f/f*^. Histological enumerations and measurements represent the mean from >10 villi per mouse. All mice were aged between 6-9 weeks. H&E = Hematoxylin and Eosin. \**p* ≤ 0.05, \*\**p* ≤ 0.01, \*\*\**p* ≤ 0.001, \*\*\*\**p* ≤ 0.001 by 2-way ANOVA with Tukey’s multiple comparisons test (E-K) or Mann Whitney test (L).

### Aberrant type 2 inflammation in *Hoil1*^−/−^ ileum is not dependent on T cells

Type 2 CD4^+^ helper T cells (Th2) and group 2 innate lymphoid cells (ILC2) are considered to be the primary producers of IL-13 and IL-4, and ILC2 are almost exclusive producers of IL-5^23^. We sought to identify the major producers of IL-13 in the ileum of *Hoil1*^−/−^ mice. First, we determined that *Il4*, *Il5* and *Il13* mRNAs were expressed primarily in the LP cell fraction (Fig. 4A). Using flow cytometry, we observed that small percentages of CD3^+^ cells (T cells) and of CD11b^+^ (myeloid) cells expressed IL-13 upon stimulation, and were similar in *Hoil1*^+/+^ and *Hoil1*^−/−^ LP (Fig. 4B, C). However, 10-15% of CD3^−^CD11b^−^CD19^−^ cells expressed IL-13, and a higher percentage of these cells were producing IL-13 in the *Hoil1*^−/−^ LP. This cell fraction includes ILCs, NK cells, dendritic cells, and mast cells. The percentage of CD3^+^ T cells was significantly reduced in *Hoil1*^−/−^ LP, indicating that these cells may also be dysregulated in the absence of HOIL1 (Fig. 4B, C).

**Figure 4:**
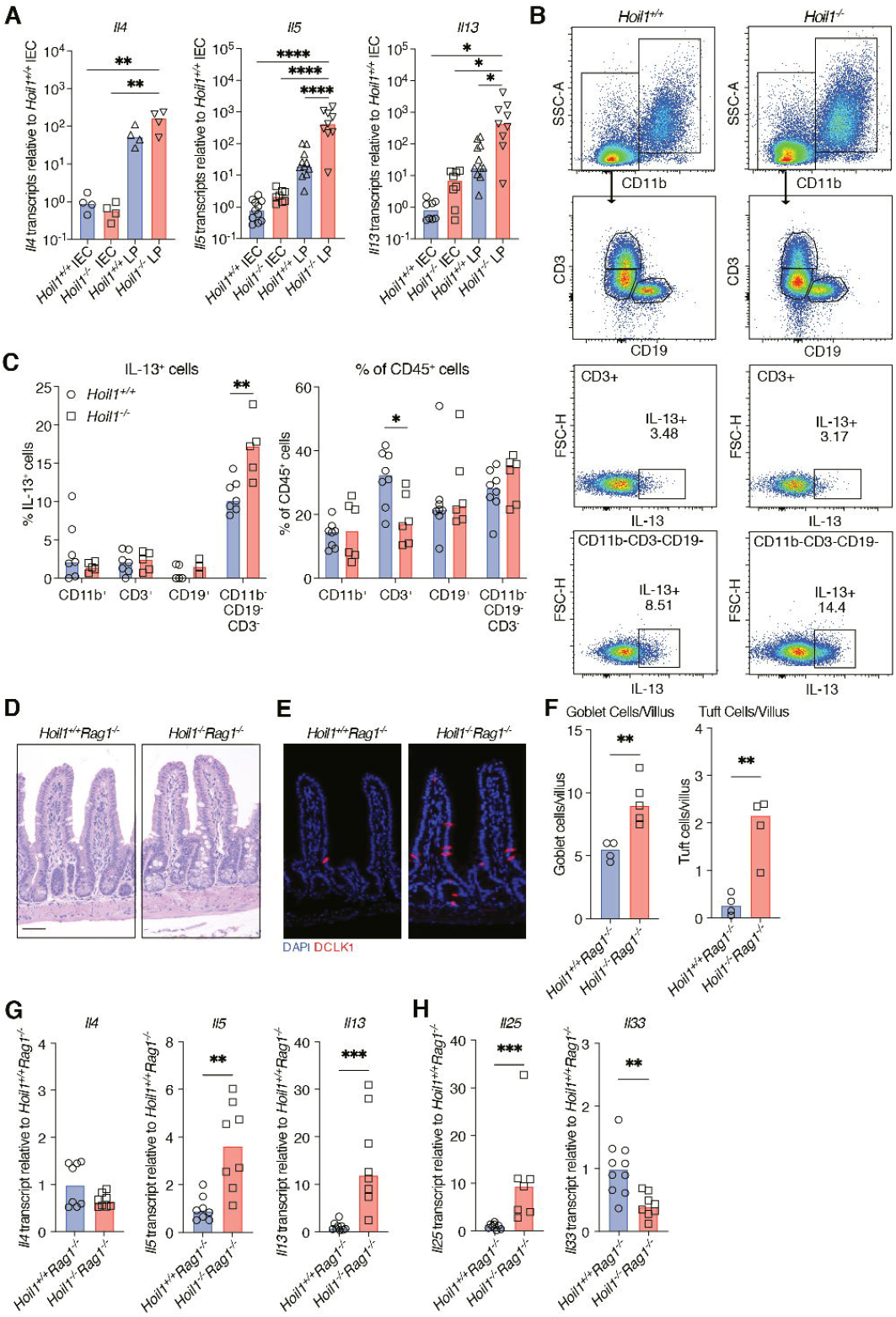
T cells are not required to drive type 2 inflammation in *Hoil1*^−/−^ ileum. (**A**) *Il4*, *Il5* and *Il13* mRNA levels in *Hoil1^+/+^* and *Hoil1*^−/−^ IEC and LP cell fractions relative to *Hoil1^+/+^* IEC. (**B**) Representative flow plots gated on live, CD45^+^ LP cells from *Hoil1^+/+^* and *Hoil1*^−/−^ ileum showing the gating strategy and intracellular IL-13 expression in CD3^+^ and in CD11b^−^ CD3^−^ CD19^−^ cell populations. (**C**) Quantification of IL-13^+^ cells (left panel) and percentage (of total CD45^+^ cells, right panel) for the indicated cell populations from *Hoil1*^+/+^ and *Hoil1*^−/−^ ileum. (**D-E**) H&E (D) and DCLK1 and DAPI (E) stained sections of ileum from *Hoil1*^+/+^*Rag1*^−/−^ and *Hoil1*^−/−^*Rag1*^−/−^ mice (scale = 50 μm). (**F**) Enumeration of goblet cells (left panel) and tuft cells (right panel) per villus in ileum from *Hoil1*^+/+^*Rag1*^−/−^ and *Hoil1*^−/−^*Rag1*^−/−^ mice. (**G-H**) *Il4*, *Il5*, *Il13* (**G**), *Il25* and *Il33* (**H**), mRNA levels in *Hoil1^+/+^Rag1^−/−^* and *Hoil1^−/−^Rag1^−/−^* distal ileum relative to *Hoil1^+/+^Rag1^−/−^*. Each symbol represents a sample from an individual mouse and bars represent the median. All mice were aged between 6-9 weeks. \**p* ≤ 0.05, \*\**p* ≤ 0.01, \*\*\**p* ≤ 0.001, \*\*\*\**p* ≤ 0.0001 by 2-way ANOVA with Tukey’s multiple comparisons test (A) or Mann-Whitney test (C, F-H).

To determine whether T cells are required for the type 2 inflammation in the absence of HOIL1, we examined *Hoil1*^−/−^*Rag1*^−/−^ mice. The ileum of *Hoil1*^−/−^*Rag1*^−/−^ mice displayed inflammatory features including increased numbers of goblet cells and tuft cells compared to *Hoil1*^+/+^*Rag*^−/−^ ileum (Fig. 4D-F). *Il5*, *Il13* and *Il25* mRNAs were elevated in *Hoil1*^−/−^*Rag1*^−/−^ compared to *Hoil1*^+/+^*Rag1*^−/−^ ileum (Fig. 4G, H), indicating that T cells are not required to trigger the inflammatory phenotype. However, *Il4* expression was not elevated in *Hoil1^−/−^Rag1^−/−^* ileum compared to *Hoil1^+/+^Rag1^−/−^* (Fig. 4G), indicating that T cells are the major producers of IL-4, as expected. Together, these data show that T cells are not required for type 2 inflammation in HOIL1-deficient ileum, and suggest that another cell type, such as ILC2, is required.

### Expression of HOIL1 is required in radiation-resistant cells to regulate intestinal type 2 inflammation

Since T cells were not required to drive type 2 inflammation caused by HOIL1-deficiency, we next questioned whether HOIL1 expression is required in cells derived from the bone marrow. We generated reciprocal bone marrow chimeras after lethal irradiation of *Hoil1^+/+^* (WT) and *Hoil1*^−/−^ (KO) mice, and chimerism was confirmed by measuring expression of *Hoil1* (*Rbck1*) mRNA in the ileum and relative amounts of *Hoil1^+/+^* and *Hoil1*^−/−^ genomic DNA in blood (Fig. 5A, B). Histological and gene expression analyses performed after 16 weeks revealed that transfer of KO bone marrow into WT mice was not sufficient to trigger goblet cell hyperplasia or excessive *Il13* expression (Fig. 5C-E). Furthermore, transfer of WT bone marrow into KO mice was not sufficient to suppress goblet cell hyperplasia or *Il13* induction. These data indicate that expression of HOIL1 in a radiation-resistant, non-bone marrow-derived cell type is required to prevent aberrant type 2 inflammation.

**Figure 5:**
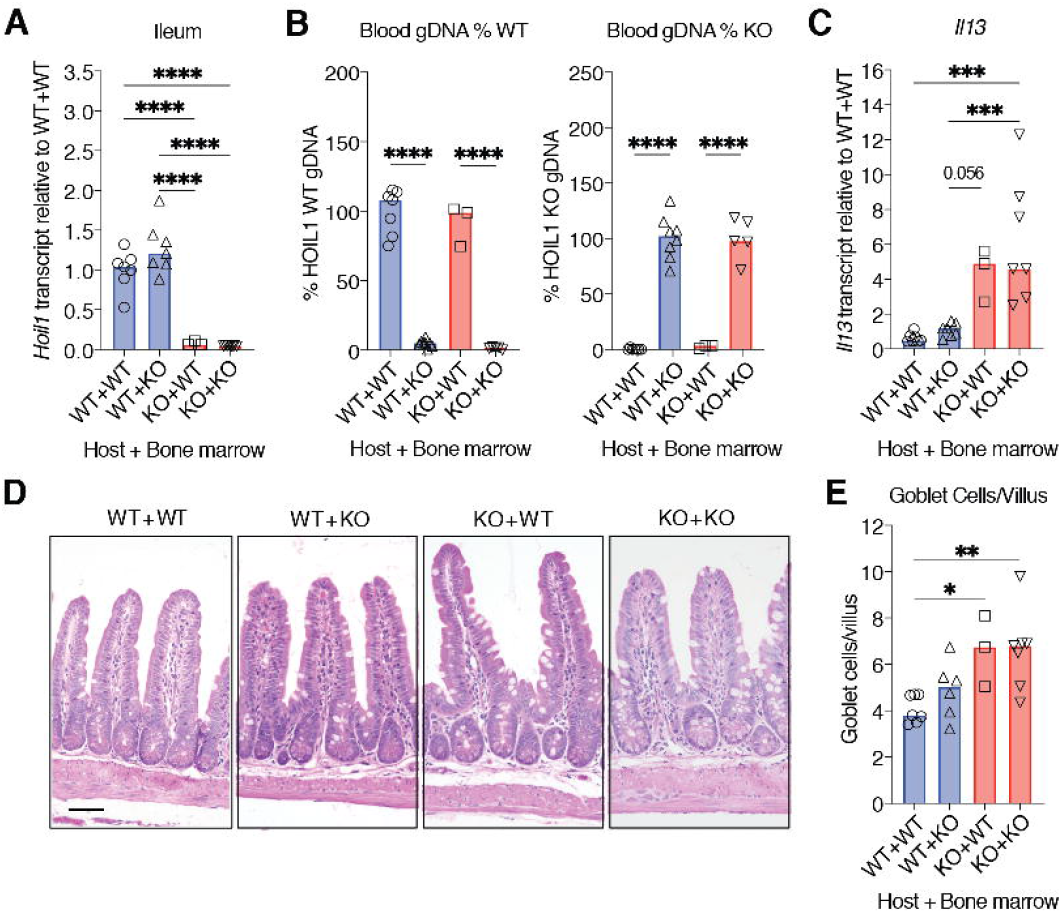
Expression of HOIL1 in non-hematopoietic cells is required to suppress type 2 inflammation in the ileum. (**A**) *Rbck1* mRNA levels (*Hoil1*, exons 3-4) in ileum from bone marrow chimeric mice relative to WT+WT. WT+WT: *Hoil1*^+/+^ mice with *Hoil1*^+/+^ bone marrow; WT+KO: *Hoil1*^+/+^ mice with *Hoil1*^−/−^ bone marrow; KO+WT: *Hoil1*^−/−^ mice with *Hoil1*^+/+^ bone marrow; and KO+KO: *Hoil1*^−/−^ mice with *Hoil1*^−/−^ bone marrow. (**B**) Percentage of WT (*Rbck1* intron 7, left panel) or KO (neomycin-resistance cassette, right panel) gDNA in blood from bone marrow chimeric mice relative to WT+WT or KO+KO controls. (**C**) Relative *Il13* mRNA levels in ileum from bone marrow chimeric mice relative to WT+WT. (**D**) Representative H&E stained sections of ileum from bone marrow chimeric mice (scale = 50 μm). (**E**) Enumeration of goblet cells per villus in ileum from bone marrow chimeric mice. Each symbol represents a sample from an individual mouse and colored bars represent the median. Histological enumerations and measurements represent the mean from >10 villi per mouse. Chimeric mice were analyzed 16 weeks after reconstitution. H&E = Hematoxylin and Eosin. \**p* ≤ 0.05, \*\**p* ≤ 0.01, \*\*\**p* ≤ 0.001, \*\*\*\**p* ≤ 0.001 by ordinary one-way ANOVA with Tukey’s multiple comparisons test.

### HOIL1 limits ILC2 numbers in the small intestine

We considered that HOIL1 may control the production of a factor that regulates type 2 cytokine expression. We previously determined that changes in expression of *Il18, Tslp*, *Il25* or *Il33* were unlikely to be responsible (Fig. 3). To assess a broader range of potential regulators, we examined the ileum from *Hoil1*^−/−^*Il4ra*^*ΔIEC*^ mice since the epithelial changes and IL-25 induction are blocked, but *Il13* and *Il5* mRNA overexpression persists in these mice (Fig. 3). We measured the mRNA expression of a number of factors that have been shown either to suppress or to promote the production of type 2 cytokines^18,19^. However, *Il10*, *Tgfb1*, *Il12p40, Tnfsf15* (*TL1A*), *Tnfsf18* (*GITRL*), *Il6, Csf2* (*GM-CSF*) *or Il1b* mRNAs were not consistently dysregulated in tissue from both *Hoil1*^−/−^*Il4ra*^*f/f*^ and *Hoil1*^−/−^*Il4ra*^*ΔIEC*^ mice relative to their HOIL1-sufficient littermates (Fig. 6A). *Il27* mRNA was significantly reduced, and IL-27 has been shown to regulate ILC and CD4^+^ T cell responses^24–26^.

**Figure 6:**
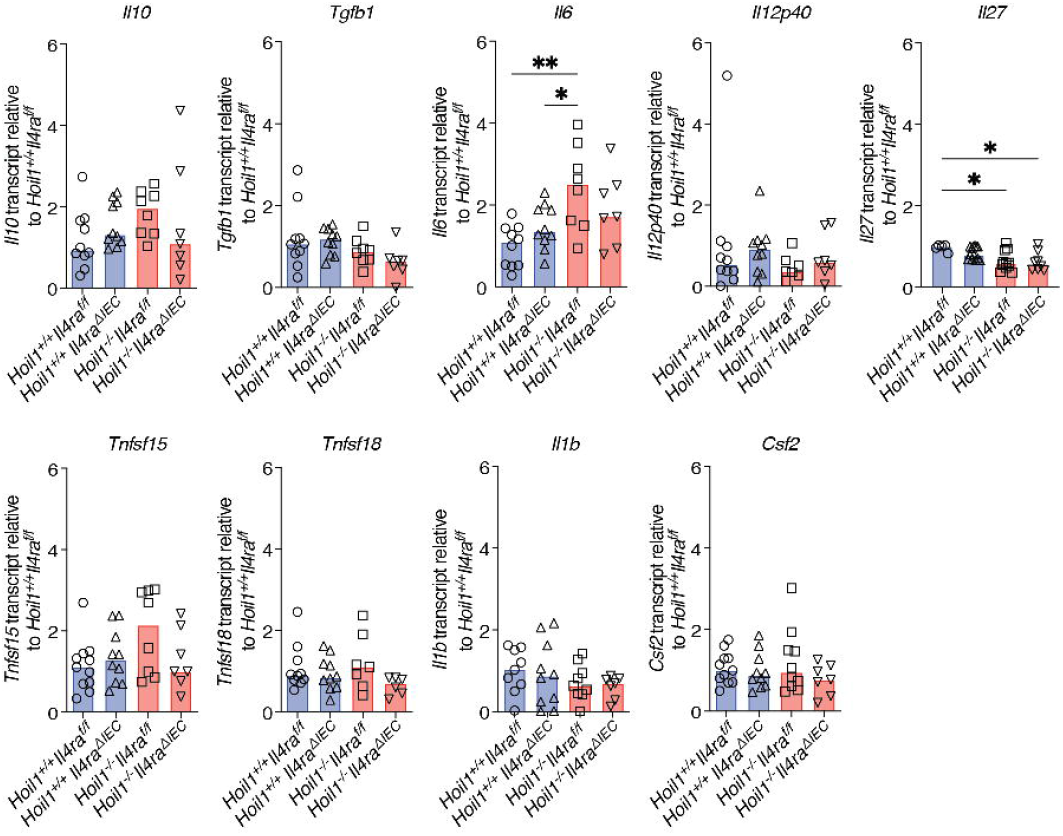
mRNA expression of common regulatory cytokines is unaltered by HOIL1-deficiency. mRNA levels for the indicated genes in distal ileum from *Hoil1*^+/+^*Il4ra*^*f/f*^, *Hoil1*^+/+^*Il4ra*^*ΔIEC*^, *Hoil1*^−/−^*Il4ra*^*f/f*^ and *Hoil1*^−/−^*Il4ra*^*ΔIEC*^ mice relative to *Hoil1*^+/+^*Il4ra*^*f/f*^. Each symbol represents a sample from an individual mouse and bars represent the median. All mice were aged between 6-9 weeks. \**p* ≤ 0.05, \*\**p* ≤ 0.01 by 2-way ANOVA with Tukey’s multiple comparisons test.

To take a more comprehensive approach to identifying transcriptional differences, we sorted CD45^+^ and CD45^−^ cells from the ileum of *Hoil1*^−/−^*Il4ra*^*ΔIEC*^ mice and performed RNA sequencing (Supplementary Table 1). In the CD45^−^ fraction, *Rbck1* (*Hoil1*) was the only gene identified as differentially expressed (DE) in the *Hoil1*^−/−^*Il4ra*^*f/f*^ and *Hoil1*^−/−^*Il4ra*^*ΔIEC*^ mice relative to the *Hoil1*^+/+^*Il4ra*^*f/f*^ and *Hoil1*^+/+^*Il4ra*^*ΔIEC*^ mice. However, in the CD45^+^ fraction, *Nmur1*, *1700061F12Rik*, *Klrg1*, *Il5*, *Il17rb*, *Epas1*, in addition to *Rbck1*, were identified as DE in *Hoil1*^+/+^ and *Hoil1*^−/−^ tissue, but unaffected by IL4Rα signaling on IECs (Fig. 7A, Supplementary Table 1).

**Figure 7:**
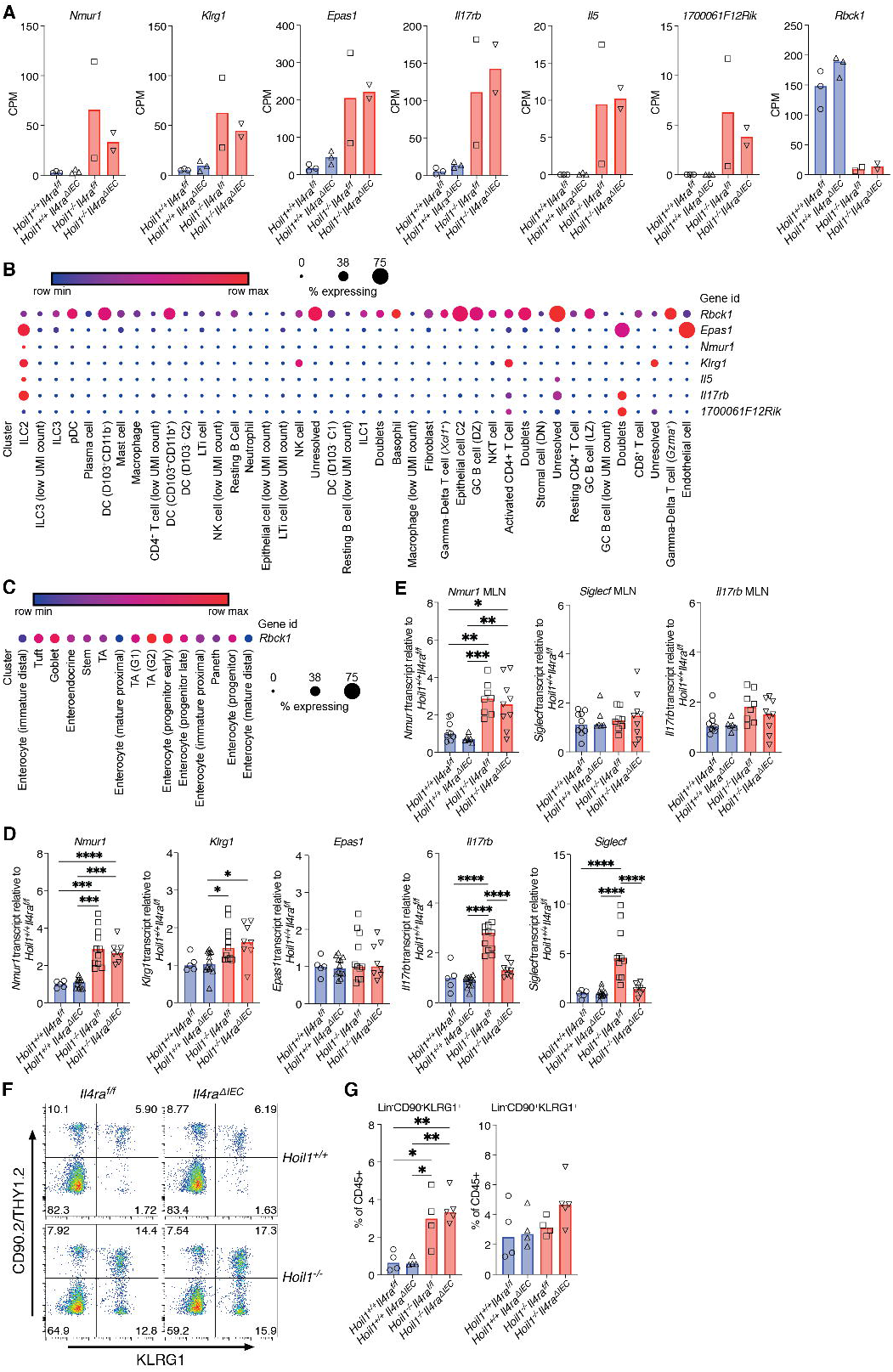
HOIL1 regulates ILC2 numbers in the small intestine. (**A**) Counts per million reads (CPM) for the indicated genes identified as differentially expressed in CD45^+^ cells from *Hoil1*^−/−^ *Il4ra*^*f/f*^ and *Hoil1*^−/−^*Il4ra*^*ΔIEC*^ ileum relative to *Hoil1*^+/+^*Il4ra*^*f/f*^ and *Hoil1*^+/+^*Il4ra*^*ΔIEC*^ by RNA-Seq analysis. *q* <0.05 for both comparisons. (**B**) Heat map/dot plot representation of the indicated genes in cell types identified in LP from untreated and OVA-treated mice by scRNAseq^27^. (**C**) Heat map/dot plot representation of *Rbck1* expression in intestinal epithelial cell subsets identified by scRNAseq^28^. (**D-E**) mRNA levels for the indicated genes in whole distal ileum (C) or MLN (D) measured by qRT-PCR and expressed as relative to *Hoil1*^+/+^*Il4ra*^*f/f*^. (**F**) Representative flow plots gated on live, CD45^+^, Lin^−^ (CD3, CD4, CD5, CD11b, CD11c, CD19, NK1.1) cells showing expression of CD90.2 and KLRG1. (**G**) Quantification of panel F for Lin^−^ CD90.2^−^ KLRG1^+^ and Lin^−^ CD90.2^+^ KLRG1+ cells (percentage of total CD45^+^ cells). Each symbol represents a sample from an individual mouse and colored bars represent the median. \**p* ≤ 0.05, \*\**p* ≤ 0.01, \*\*\**p* ≤ 0.001, \*\*\*\**p* ≤ 0.0001 by 2-way ANOVA with Tukey’s multiple comparisons test.

To determine which specific cell types express these DE genes in the small intestine during a type 2 inflammatory response, we queried the Immunological Genome Project (ImmGen) database and published single-cell RNA sequencing datasets^27,28^. *Klrg1* can be expressed by ILC2, NK cells and Th2 cells, and *Il17rb*, *1700061F12Rik* and *Epas1* can be induced in activated ILC2 and Th2 cells. However, *Il5* and *Nmur1* mRNAs are relatively specific to ILC2 among CD45+ cells within the small intestine^27,29^ (Fig. 7B). *Rbck1* expression was variable among immune cell and epithelial cell populations, with notable expression in several dendritic cell subsets, γδT cells and germinal center B cells, as well as in tuft cells, goblet cells, transit amplifying cells and enterocyte progenitors^28^ (Fig. 7B, C).

Using qRT-PCR, we confirmed that *Nmur1* was more highly expressed in distal ileum from *Hoil1*^−/−^*Il4ra*^*f/f*^ and *Hoil1*^−/−^*Il4ra*^*ΔIEC*^ mice (Fig. 7D). *Klrg1* mRNA was also slightly elevated, although this was not significant in the *Hoil1*^−/−^*Il4ra*^*ΔIEC*^ tissue. *Epas1* mRNA expression was not detectably different in whole tissue. *Il17rb* and *Siglecf* (included as a marker for eosinophils) were elevated in tissue from *Hoil1*^−/−^*Il4ra*^*f/f*^ but not *Hoil1*^−/−^*Il4ra*^*ΔIEC*^ mice, indicating that changes in their expression are dependent on IL4Rα signaling in IECs. Furthermore, *Nmur1* was more highly expressed in the MLN from *Hoil1*^−/−^*Il4ra*^*f/f*^ and *Hoil1*^−/−^*Il4ra*^*ΔIEC*^ mice, but *Siglecf* and *Il17rb* were not (Fig. 7E). NMUR1 is a neuropeptide receptor that has recently been shown to be preferentially expressed on ILC2, and to induce ILC2 activation and proliferation in response to neuromedin U (NMU) produced by mucosal neurons^30–32^. These findings suggested that ILC2 numbers or activation state are dysregulated in the small intestine of HOIL1-deficient mice. Flow cytometry revealed that the frequency of Lin^-^KLRG1^+^CD90.2^lo^ ILC2 was approximately five-fold higher in the ileum of *Hoil1*^−/−^ mice, and remained elevated even in the absence of IL4Rα signaling on IECs (Fig. 7F, G), indicating that HOIL1 limits number of ILC2 in the small intestine.

## Discussion

In this study, we have identified a critical role for HOIL1 in regulating type 2 inflammation in the small intestine of mice. HOIL1-mutant mice exhibited characteristic goblet and tuft cell hyperplasia that was associated with increased expression of IL-4, IL-5, IL-13 and IL-25, and was dependent on signaling through IL4Rα on IECs. *Il13* and *Il5* remained elevated when tuft cell and IL-25 induction were blocked, demonstrating that HOIL1 functions upstream of IL4Rα in the feed-forward cycle.

Although Th2 cells, ILC2, eosinophils and mast cells can express type 2 inflammatory cytokines, analysis of *Hoil1*^−/−^*Rag1*^−/−^ mice demonstrated that T cells are not required for inflammation. These findings were consistent with an increase in intracellular IL-13 observed in a CD11b^−^CD3^−^CD19^−^ cell population, but not in the CD3^+^ or CD11b^+^ cell populations from the HOIL1-deficient small intestine. Furthermore, RNA-Seq analysis of CD45^+^ cells identified an increase in mRNA expression of six ILC2-associated genes, two of which (*Nmur1* and *Il5*) are specific for ILC2. Subsequent flow cytometric analysis revealed a four to five-fold increase in Lin^-^KLRG1^+^CD90^lo^ ILC2 in HOIL1-deficient tissue, which was independent of IL4Rα signaling on IEC and induction of IL-25. These KLRG1^+^CD90^lo^ ILC2 may be similar to the inflammatory ILC2 that have been reported to proliferate in the small intestine, then migrate to the lung and other tissues in response to helminth infection or IL-25 treatment^33,34^.

The proliferation and activation of ILC2 can be induced by IL-25, TSLP and IL-33, along with additional signals such as cysteinyl leukotrienes, NMU, or Notch ligands^30–32,35–37^. We were unable to detect differences in TSLP or IL-33 mRNA or protein expression and, although IL-25 was elevated in HOIL1-deficient ileum, the increase in IL-25 was not required. Global mRNA analysis of CD45^+^ and CD45^-^ cells did not reveal candidates except genes associated with ILC2. One possibility is that HOIL1 plays a cell-intrinsic role in regulating ILC2, and this would be consistent with a requirement for HOIL1 in radiation-resistant, non-bone marrow derived cells, since some ILC2 are thought to self-renew in tissues^38^. A recent study identified LUBAC as a component of the IL17RA/IL17RC receptor signaling complex (RSC) required for efficient signal transduction and NFκB activation^39^. The same study identified a negative feedback loop for the IL-17RSC, although LUBAC did not appear to be involved. Since IL-25 signals though IL17RB, which is highly expressed on gut ILC2, it is plausible that HOIL1 and LUBAC regulate tonic IL-25/IL17RB signaling and therefore ILC2 numbers and activation state (Fig. 8). Other mechanisms are possible including ILC2-intrinsic regulation of IL17RB expression or signaling through other receptors, or ILC2-extrinsic roles for IL-27, interferons, neuropeptides such as NMU, or lipid mediators such as prostaglandins and leukotrienes. Further studies, such as cell type-specific deletion of *Hoil1*, will be required to distinguish these possibilities.

**Figure 8:**
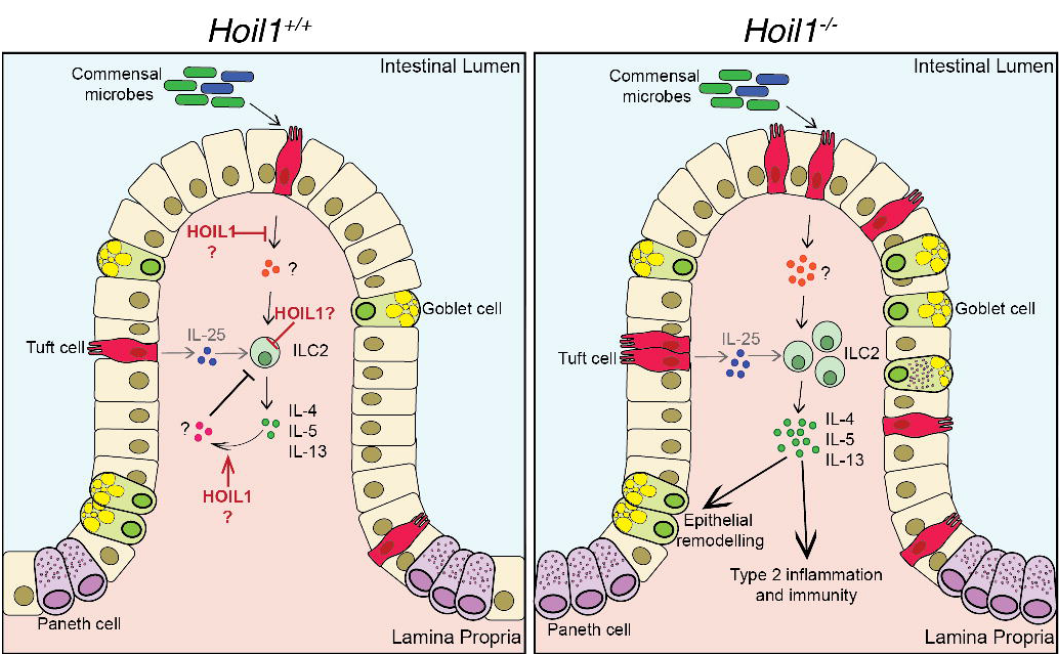
Model of role of HOIL1 in regulating type 2 inflammatory signaling. HOIL1 acts in a radiation-resistant cell type to suppress ILC2 proliferation and the production of IL-4, IL-5 and IL-13 in the presence of commensal microbes. HOIL1 may function to suppress the production of a positive regulatory factor (orange circles) upstream of IL-4, IL-5 and IL-13, or may be required for the negative regulation of ILC2 through a cell-extrinsic (pink circles) or cell-intrinsic mechanism, such as inhibition of IL17RB (IL25R) signaling. HOIL1 functions outside of the IL-13 – tuft cell – IL-25 feed-forward loop. In the absence of HOIL1, excessive IL-4, IL-5 and IL-13 can trigger chronic type 2 inflammation including goblet and tuft cell hyperplasia.

Although IL-25 is a well-established activator of intestinal ILC2, examination of antibiotics-treated mice indicated that a signal other than IL-25 is involved. Antibiotics treatment reduced *Il4*, *Il5* and *Il13* mRNA levels in *Hoil1*^−/−^ tissue to levels similar to *Hoil1*^+/+^ tissue from water-treated mice. *Il25* mRNA, however, was only partially reduced by antibiotics, and reduction of *Il25* mRNA (by blocking IL4Rα signaling) was not sufficient to reduce *Il5* and *Il13* mRNA. Others have shown that resting ILC2 numbers and *Il5* expression are largely unaffected by the absence of microbes in wild-type germ-free mice^29^. However, the additional ILC2 we observed in the absence of HOIL1 may be activated ILC2 (CD90^lo^ or CD90^−^)^33,34^ and subject to additional modes of regulation. Future studies will need to determine whether loss of microbial exposure reduces ILC2 numbers or activation state, and to determine whether HOIL1 regulates this response to microbes in an ILC2-intrinsic or extrinsic manner (Fig. 8). Analysis of scRNA-seq datasets revealed that *Hoil1* is expressed in a variety of cell types with notable expression in tuft cells, goblet cells and transit amplifying cells that could sense microbial products. We did not identify any differentially expressed genes among the CD45^-^ cells, although this may be due to the limited power of the analysis, or to non-transcriptional regulation of key molecules. *Hoil1* was also highly expressed in several dendritic cell and B cell subsets that may warrant further analysis. Since HOIL1 has recently been shown to be a functional E3 ubiquitin ligase^40,41^, it will be important to determine whether linear ubiquitination by HOIP or the E3 ligase activity of HOIL1 regulates type 2 inflammatory responses in the small intestine.

Type 2 responses are critical for immune responses against extracellular parasites such as intestinal worms, yet aberrant type 2 inflammation drives allergy, asthma and atopic dermatitis. We have identified HOIL1 as an important regulator of intestinal ILC2 and type 2 inflammation, contributing to our broader understanding of the mechanisms regulating intestinal homeostasis and inflammation.

## Methods

### Mice

All mice used in this study were on a C57BL/6J background. Mice were housed and bred in accordance with Federal and University guidelines and protocols were approved by the University of Illinois at Chicago Animal Care Committee and the Animal Studies Committee of Washington University. *Hoil1*^−/−^ (Rbck1^tm1Kiwa^) and *Hoil1*^−/−^*Rag1*^−/−^ mice have been described previously^8,13,21^. Co-housed *Hoil1*^+/+^ littermates were used as wild type controls. *Il4ra*^*flox/flox*^ mice^21^ were a gift from Ajay Chawla and bred to *Hoil1*^−/−^ mice. VillinCre (B6.Cg-Tg(Vil1-cre)997Gum/J) mice were purchased from The Jackson Laboratory and bred to *Hoil1*^−/−^*Il4ra*^*flox/flox*^ mice. Both male and female mice were included in all analyses. Mice from at least two litters were used to generate each data set, providing 3 to 10 mice per group. No mice were excluded from the analyses.

### Intestinal epithelial cell and lamina propria cell separation for mRNA analysis

Whole ileum and jejunum was flushed with PBS, the Peyer’s patches removed, opened longitudinally, and cut into 1 cm pieces. Two consecutive washes were performed with HBSS supplemented with 10% bovine calf serum, 15 mM HEPES, 5mM EDTA and 1.25 mM DTT at room temperature for 20 min under continuous rotation followed by 20 seconds of vortexing in PBS pH 7.4. IECs were collected and resuspended in TRI-reagent (Sigma). Remaining tissue containing the lamina propria cell fraction was homogenized in TRI-reagent as described below.

### Ileum digestion

Whole ileum and jejunum was prepared as described above for epithelial cell and lamina propria (LP) separation. The remaining LP pieces were transferred to 15 ml RPMI 1640 supplemented with 10% FBS, 50 U/ml penicillin, 50 μg/ml streptomycin and 2 mM L-glutamine with 0.5 mg/ml Collagenase VIII (Sigma). Samples were shaken vigorously by hand and placed in a shaking incubator at 220 rpm and 37°C for 15 min with additional manual shaking at 10 min. After 15 min, 35 ml ice-cold complete media. Samples were washed twice with FACS buffer (PBS pH 7.4 plus 1% FBS and 2 mM EDTA) and filtered with 100 μm and 70 μm strainers.

### Flow cytometry

Single cell suspensions were incubated with CD16/CD32, normal mouse, rat and hamster serum to block non-specific binding, then stained with fluorophore-conjugated antibodies against: CD45 (30-F11), CD11b (M1/70), CD19 (6D5), Siglec-F (S17007L), CD3 (17A2), IL7R-α (A7R34). Viable cells were identified by exclusion of Zombie NIR fixable viability dye (BioLegend). For identification of ILC2, lineage-positive cells were excluded by staining for CD3 (17A2), CD4 (GK1.5), CD5 (53-7.3), CD11b (M1/70), CD11c (N418), CD19 (6D5) and NK1.1 (PK136). CD45 (30-F11), CD90.2 (30-H12), and KLRG1 (2F1) were used for positive identification. For intracellular staining, cells were re-stimulated for 4 hr with PMA/Ionomycin and Brefeldin A (Biolegend), according to manufacturer’s recommendations, then fixed and permeabilized using a Foxp3/Transcription Factor buffer set (eBioscience), according to manufacturer’s instructions. Cells were stained with a fluorophore-conjugated antibody against IL-13 (eBio13A). Flow cytometry was performed using a BD Fortessa X20 or CytoFLEX S (Beckman Coulter), and data were analyzed using FlowJo10.5 (TreeStar Inc.). Flow cytometry gating strategy was based on fluorescence minus one (FMO), unstained, and isotype controls.

### Fluorescence-activated cell sorting and RNA-Seq analysis

IEC and LP cell fractions were prepared as described above, and 50% of the IEC combined with the LP cells. Cells were stained with Zombie NIR fixable viability dye (BioLegend) and AF488 anti-CD45 (30-F11). Live, CD45^+^ and CD45^−^ singlets were sorted on a MoFlo Astrios Cell Sorter (Beckman Coulter) into RLT buffer containing 1% β-mercaptoethanol, flash frozen on dry ice and stored at −80°C. Three samples per group were generated, for a total of 24 samples.

Total RNA was purified using an RNeasy Plus Micro Kit (Qiagen) according to the manufacturer’s instructions. Libraries for Illumina sequencing were prepared using a Universal Plus mRNA-Seq library preparation kit (Tecan/NuGen, 0520-A01), using 10-20 ng RNA per sample. In brief, RNA underwent poly-A selection, enzymatic fragmentation, and generation of double-stranded cDNA using a mixture of oligo (dT) and random priming. The cDNA underwent end repair, ligation of dual-index adaptors, strand selection, and 16 cycles of PCR amplification. All purification steps were carried out using Agencourt AMPure XP Beads (Beckman Coulter A63881). Concentration of the final library pool was confirmed by qPCR and subjected to test sequencing in order to check sequencing efficiencies and adjust proportions of individual libraries accordingly. The pool was purified with the Agencourt AMPure XP Beads, and sequenced on a NovaSeq 6000 S4 flow cell, 2 x 150 bp, approximately 30 M clusters per sample, at the University of Illinois Roy J. Carver Biotechnology Center High-Throughput Sequencing and Genotyping Unit.

Raw reads were trimmed to remove adapters and bases from the 3’ end with quality scores less than 20 using cutadapt; trimmed reads shorter than 40bp were discarded. Reads were aligned to the mouse reference genome (mm10) in a splice-aware manner using the STAR aligner^42^. ENSEMBL gene and transcript annotations were used, which included non-coding RNAs in addition to mRNAs. The expression level of each gene was quantified using FeatureCounts^43^ first as raw read counts, which were suitable for differential expression analyses, and also normalized to counts-per-million for direct comparison between samples.

Differential expression statistics (fold-change and *p*-value) were computed using edgeR^44,45^ on raw expression counts obtained from quantification, using the generalized linear model framework to test for effects due to *Hoil1* and *Il4ra* expression simultaneously, including interaction between those factors, in addition to pair-wise comparisons between different groups using exactTest. Analyses were performed separately for CD45^+^ and CD45^−^ cell samples. Three outlier samples, identified by principal component analysis, were removed from the analyses. *p*-values were adjusted for multiple testing using the false discovery rate (FDR, *q*-value) correction of Benjamini and Hochberg. Differentially expression genes were determined based on an FDR threshold of 5% (0.05) in the multi-group comparison.

### Analysis of scRNA-seq datasets

scRNA-seq datasets of small intestine LP (GEO: GSE124880)^27^ and epithelium (GEO: GSE92332)^28^ were probed for expression of our genes of interest through the Broad Institute’s Single Cell Portal.

### Antibiotic treatment

Mice were treated by daily oral gavage with either sterile dH_2_O or 100 mg/kg ampicillin, 100 mg/kg neomycin, 50 mg/kg vancomycin, and 100 mg/kg metronidazole dissolved in sterile dH_2_O^46^. Body weight was monitored daily, and stool pellets collected on days 0, 4, 7 and 14. Two independent experiments were performed. Randomization of animals into treatment groups was not explicitly performed, but determined by cage assignment at weaning prior to genotyping.

### Fecal DNA isolation

DNA was isolated from homogenized fecal pellets by double phenol:chloroform:isoamyl alcohol extraction and isopropanol precipitation^47^.

### Tritrichomonas muris testing

Fecal pellets were collected from at least two breeding cages from each mouse strain: *Hoil1*^+/−^, *Hoil1*^+/−^*Rag1*^−/−^ *Hoil1*^+/−^*l4ra*^*f/f*^*Vil1cre* (2-3 pellets per cage), and shipped to IDEXX BioAnalytics for testing for *Tritrichomonas muris*. All samples were negative.

### RNA isolation

Whole 1 cm tissue samples of distal ileum (1 cm from the cecum), jejunum (10-11 cm from the stomach), distal colon, or mesenteric lymph nodes were snap-frozen and stored at −80°C. Frozen samples were homogenized in TRI-Reagent (Sigma) using zirconia/silica beads (BioSpec) and a Mini-Beadbeater 24 (BioSpec). RNA was isolated according to the manufacturer’s instructions. RNA samples were treated with Turbo DNA-free DNase (Invitrogen) and 1 μg of RNA used as a template for cDNA synthesis with random primers and ImProm-II reverse transcriptase (Promega).

### Quantitative PCR

Quantitative PCR (qPCR) was performed on a QuantStudio 3 Real-Time PCR System (Applied Biosytems) using predesigned 5’ nuclease probe-based assays for: *Il4* (Mm.PT.58.32703659), *Il5* (Mm.PT.58.41498972), *Il13* (Mm.PT.58.31366752), *Ifng* (Mm.PT.58.41769240), *Tnf* (Mm.PT.58.12575861), *Il18* (Mm.PT.58.42776691), *Il25* (Mm.PT.58.28942186), *Il33* (Mm.PT.58.12022572), *Tslp* (Mm.PT.58.41321689), *Rbck1* (Mm.PT.58.30767649), *Il6* (Mm.PT.58.10005566), *Il10* (Mm.PT.58.13531087), *Il12p40* (Mm.PT.58.12409997), *Tgfb* (Mm.PT.58.11254750), *Tnfsf15* (Mm.PT.58.43933933), *Tnfsf18* (Mm.PT.56a.8500128), *Csf2* (Mm.PT.58.9186111), *Il1b* (Mm.PT.58.41616450), *Il27* (p28, Mm.PT.58.11487953), *Nmur1* (Mm.PT.58.32232111), *Klrg1* (Mm.PT.58.30803964), *Epas1* (Mm.PT.58.13819524), *Il17rb* (Mm.PT.58.12616779), *Siglecf* (Mm.PT.58.6685529) (Integrated DNA Technologies). 16s qPCR was performed using PowerSYBR Green assay (Invitrogen) and primers: 515F (5’-GTG CCAGCMGCCGCGGTAA-3’) and 805R (5’-GACTACCAGGGTATCTAATCC-3’)^47^. Transcript levels were quantified using the relative standard curve method, with *Rps29* as the reference gene^7^. qPCR standards were designed using assay primer sequences and assembled in gene blocks (Integrated DNA Technologies).

### Histology

Distal ileum (last 6 cm of the small intestine up to the cecum) was flushed with PBS followed by 10% buffered formalin and opened longitudinally, flattened and pinned in 10% buffered formalin for 24 hr followed by washing and dehydration with 70% ethanol. Strips of tissue were embedded in 2% agar prior to paraffin embedding. Blocks were sectioned and stained with Hematoxylin solution Gill III (MilliporeSigma) and 1% Eosin Y Solution (G-Biosciences). For immunofluorescent staining, antigen retrieval was performed by boiling in 10 mM Tri-sodium citrate (dihydrate) with 0.05% Tween 20, pH 6.0 for 20 min. Sections were blocked with PBS containing 5% FBS and 0.1% TRITON X-100 for 3 hr, and then incubated with rabbit anti-mouse DCLK1 (ab31704, Abcam) or rabbit anti-mouse Lysozyme (ab108508, Ambcam) in PBS + 5% FBS at 4°C overnight. Sections were washed with PBS + 0.3% Tween20 and incubated with Alexa Fluor 555 donkey anti-rabbit (A31572, Invitrogen) for 1 hr at 4°C. Sections were washed with PBS + 0.3% Tween20, and counterstained and mounted with Prolong Gold antifade reagent with DAPI (Invitrogen). Imaging was performed on a BZ-X710 microscope (Keyence). Tuft and goblet cell quantification was based on an average of at least 10 villi and crypts per mouse. Blinding was performed by assigning slides a mouse tag number, and matching to genotype post-quantification.

### Bone marrow chimeric mice

Recipient mice were exposed to 1200 rad of whole body irradiation and injected intravenously with 10 million whole bone marrow cells from donor mice. Mice were allowed to reconstitute for 16 weeks before sacrifice for analysis of intestinal tissue. Mice were bled at 12 to 14 weeks post-irradiation to determine percent chimerism. Genomic DNA was isolated from peripheral blood and analyzed by qPCR for the presence of *Rbck1/Hoil1* intron 7 (in control cells; 5′-ATG CTGGAGTAGAGGCTGGA-3′ and 5′-TGACTGCTGCTTGGAGAGTG-3′), or the neomycin-resistance cassette (in *Hoil1*^−/−^ cells; 5′-CAAGATGGATTGCACGCAGG-3′ and 5′-GCAGCCGAT TGTCTGTTGTG-3′). *Rag2* was used as a normalization control (5′-GGGAGGACACTCACTTGCCAGTA-3′ and 5′-AGTCAGGAGTCTCCATCTCACTGA-3′).

### Total protein isolation and ELISAs

Whole 1 cm tissue samples of distal ileum (1 cm from the cecum) were homogenized in PBS with Halt phosphatase and protease inhibitors (Thermo scientific) using sterile zirconia/silica beads (BioSpec) and a Mini-Beadbeater 24 (BioSpec). Supernatant was reserved for further analysis and total protein quantified using DC Protein assay (Bio-Rad). Cytokine production was determined in distal Ileum by ELISA using R&D DuoSet for IL-33 and IL-13, and Biolegend ELISA MAX for TSLP, IL-4, IL-5, and IL-25 following the manufacturers’ instructions and analyzed with a microplate reader (BioTek Synergy 2).

### Statistical analyses

Data were analyzed with Prism 9 software (GraphPad Software, San Diego, CA). Statistical significance was determined by tests as indicated in the figure legends.

## Data availability

RNA-Seq data are available at NCBI Gene Expression Omnibus (GEO) through accession number GSE196550.

## Acknowledgements

We would like to acknowledge D. Kreamalmeyer, Joy Loh, Christine T. Luo, X. Zhang, M. Byrne, B. Studnicka, A. Darbandi, the Research Histology and Tissue Imaging Core, and the Flow Cytometry Core at the University of Illinois Chicago, and the Digestive Diseases Research Core Center and the Developmental Biology Histology Core at Washington University for technical assistance. RNA sequencing was performed by the UIC Genome Research Core, and bioinformatics analysis was performed by the UIC Research Informatics Core, supported in part by NCATS through Grant UL1TR002003. We thank Skip Virgin and members of the MacDuff laboratory for helpful discussions. We would like to thank A. Chawla for providing the *Il4ra*^*flox/flox*^ mice. This work was funded by UIC institutional start-up funds to DAM.

## Author Contributions

MJW, TCL, KI, TSS and DAM designed the study; MJW, JNM, VLH and DAM performed experiments; MJW, JNM, VLH and DAM analyzed the data and performed statistical analyses; MJW, VLH, TCL, TSS and DAM interpreted the data; MJW and DAM drafted the manuscript; all authors were involved in discussing the data and provided feedback on the manuscript.

